# Biosynthetic gene clusters everywhere, but the environment selects

**DOI:** 10.64898/2026.01.28.702283

**Authors:** Zander Rainier Human, Rene Hoover, Clara Chan, Kirsten Küsel, Carl-Eric Wegner

## Abstract

The *in situ* relevance of biosynthetic gene clusters (BGCs) remains poorly understood. We applied meta-omics to characterize BGC diversity and activity along a peatland redox gradient. From seven metagenomes, we recovered 9,694 BGCs spanning diverse taxa, most lacking close relatives in reference databases, indicating extensive novelty. Only 9-27% of this potential was expressed in situ, with Acidobacteriota, despite moderate repertoires, accounting for over half of all BGC transcription. “Talented producers” with up to 24 clusters were largely silent, and expression was inversely related to BGCs per genome. Acidobacteriota and other oligotrophic taxa expressed most of their BGCs, whereas copiotrophic Pseudomonadota expressed a few, suggesting that life-history and stress-responsive regulation govern activation. PKS and terpene BGCs expression was enriched in genomes expressing markers of aerobic respiration and stress, whereas RiPPs were associated with antimicrobial resistance, stationary phase and stress. Thus, although BGCs are widespread, environmental constraints determine which are realized.

## Introduction

Secondary metabolites are synthesized in an assembly-line fashion by enzymes encoded by biosynthetic gene clusters (BGCs), which are groups of colocalized and coregulated genes. These metabolites are functionally diverse and include metal chelators, antifungals, and antibiotics^1–5^. The biosynthesis of secondary metabolites is energetically costly, and most known producers are aerobes^6^. This has contributed to the view that secondary metabolism is favored in energy-rich, oxic environments. At the same time, environmental triggers and physiological transitions often provide the cues that activate secondary metabolite production in culture, including nutrient limitation^7,8^, especially nitrogen and phosphorus depletion^9^, and oxidative stress^10^. In *Streptomyces*, where most BGCs remain silent under standard growth conditions, biosynthesis is commonly induced by nutrient depletion, the quality of carbon sources, entry into the stationary phase, or cell-to-cell interactions^11,12^. Similar regulatory patterns are observed in other soil bacteria, such as *Burkholderia* and *Pseudomonas*, which produce secondary metabolites typically during stationary phase or under nutrient limitation^13^. In contrast, some myxobacteria, such as *Sorangium* spp., express nonribosomal peptide synthetase (NRPS) and polyketide synthase (PKS) clusters during exponential growth^14^. Secondary metabolism is tightly coupled to environmental and physiological cues. The triggers can differ substantially across microbial groups.

The study of BGCs has historically been limited to organisms that can be readily cultivated, resulting in a focus on a restricted set of lineages, primarily Actinobacteria (Actinomycetota) and Proteobacteria (Pseudomonadota). Cultivation, however, represents a strong bottleneck: it excludes most environmental microbes, masks ecological drivers of secondary metabolism, and often fails to activate many BGCs under artificial growth conditions. Consequently, the diversity and ecological relevance of microbial secondary metabolism remain poorly understood.

Advances in metagenomics, particularly the reconstruction of metagenome-assembled genomes (MAGs), have expanded the microbial tree of life^15–17^ and uncovered vast BGC diversity across environments ranging from oceans to soils^18–21^. These surveys demonstrate that BGCs are nearly ubiquitous across microbial lineages, including those uncultured and taxonomically underrepresented. They also suggest that the synthesis and utilization of secondary metabolites strongly depend on the environment.

Peatlands represent a particularly promising but still underexplored ecosystem in this context. They are globally significant carbon reservoirs^22^ and harbor microbial communities that are phylogenetically diverse, taxonomically novel, and underrepresented in culture collections^23^. Many of these lineages, including Acidobacteriota, Verrucomicrobiota, Planctomycetota, and other poorly studied groups, have been shown to harbor rich biosynthetic potential, yet the ecological roles of these metabolites remain largely unknown^18,24–26^. Peatland hydrology creates steep vertical redox gradients, with oxic surface layers transitioning into permanently anoxic, water-saturated horizons, accompanied by shifts in nutrient availability and electron acceptor competition^27–29^. These gradients create environmental conditions that parallel the nutrient and redox cues known to induce secondary metabolism in cultured bacteria^9^.

We hypothesize that peatlands are not only reservoirs of biosynthetic novelty but also natural laboratories for testing how environmental heterogeneity filters from the broad repertoire of BGCs present in microbial genomes. Here, we test if BGCs, although widespread across peatland microbial lineages, including many beyond the classical producers, are selectively structured by redox conditions in terms of distribution and expression. By combining metagenomic and transcriptomic analyses, we assess both the diversity and *in situ* expression of BGCs along natural redox gradients and show how the environment selects from the universal biosynthetic potential of microbes.

## Results

### Metagenomes from an acidic, Fe-rich peatland

To capture microbial diversity and biosynthetic activity across the natural redox gradient of the Schlöppnerbrunnen fen, we performed genome-resolved metagenomic and metatranscriptomic analyses on peat collected between 5 and 30 cm depth. This interval spans the transition from the oxic surface layer to the anoxic, water-saturated layer, where moisture and microbial activity remain high. Samples were collected in 5 cm increments, with a second core taken 1 m away for replication.

Coverage estimations based on k-mer profiles confirmed that our deep sequencing adequately captured the genomic diversity of the peatland microbiome **(Supplementary Table 1)**. Across all depths, the peatland microbiome was dominated by members of the Acidobacteriota (43-30%), Pseudomonadota (16-19%), Actinomycetota (3-10%), Verrucomicrobiota (2.7-6.5%), and Patescibacteriota (5-11%) (**Fig. 1a, Supplementary Table 2**). The relative abundance of these phyla shifted markedly with depth. Acidobacteriota, Actinomycetota, and Verrucomicrobiota decreased in deeper layers, whereas Desulfobacterota, Archaea, and Pseudomonadota increased in abundance toward more anoxic conditions. This stratification reflects a transition from predominantly aerobic to anaerobic microbial assemblages along the redox gradient. Taxonomic composition was inferred from the abundance of single-copy housekeeping genes identified using SingleM^30^.

**Fig. 1.**
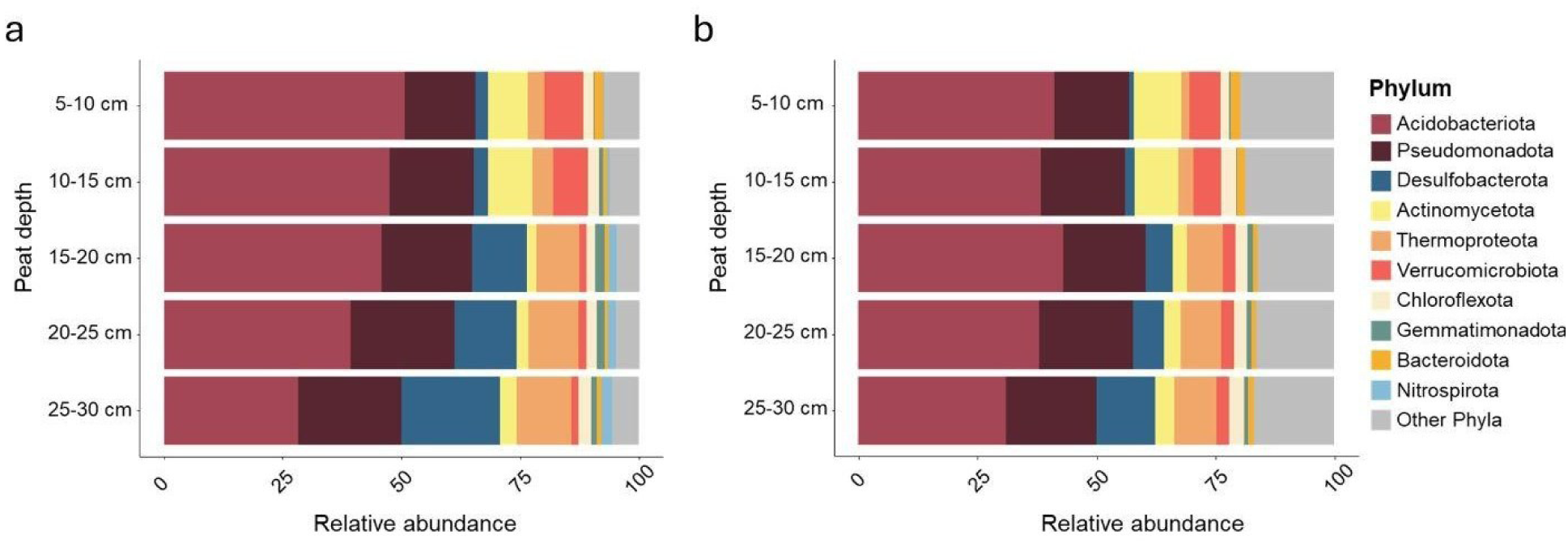
Microbial community composition and diversity across peatland depths. (a) Taxonomic profiles derived from raw reads assigned to single-copy marker genes using SingleM and (b) relative abundances of reconstructed MAGs, both shown as mean community composition across five depth intervals (5 - 30 cm).

Metagenome assemblies were clustered into 573 metagenome-assembled genomes (MAGs) with at least 70% completeness and contamination less than 10% **(Supplementary data 1)**, exceeding the MIMAG completeness standard for medium quality MAGs^31^. The majority of MAGs were assigned Acidobacteriota (131), Pseudomonadota (119 MAGs), Actinomycetota (44), Verrucomicrobiota (37), and 43 from subgroups of Desulfobacterota. From the 52 Archaea MAGs, most belonged to the Thermoproteota (31) and Halobacteriota (11) (**Supplementary table 3**).

Relative MAG abundances, determined by mapping quality-controlled reads to MAGs with coverM ^32^, were consistent with taxonomic profiles obtained from raw reads using SingleM, which assigns taxonomy based on single-copy marker genes (**Fig. 1b**, **Supplementary Table 3**). To assess how metabolic strategies varied along the redox gradient, MAG abundances were combined with O_2_-utilizing phenotype assignments predicted from genome-wide amino-acid k-mer composition^33^, and the number of oxygen-utilizing enzymes ^34^. Both strategies indicated a clear increase in anaerobic taxa with depth, from 6 to 34% in the oxic surface layer to 29% to 50% in the deepest peat layer (25 – 30 cm depth) (**Supplementary Fig. 2**).

### Identification and abundance of BGCs

The biosynthetic potential of the peatland was assessed by identifying biosynthetic gene clusters (BGCs) from contigs longer than 5 kbp using antiSMASH^35^. A total of 9,694 BGCs (**Supplementary Table 4, Supplementary data 2**) were identified, 1,594 (16.4%) were considered complete. The average length was 31,073 bp, the longest BGC was a 100,738 bp NRPS-Type I PKS-transAT-PKS-like cluster, and the shortest a 10,219 bp RiPP-like BGC.

BGCs were dereplicated into 4,669 gene cluster families (GCFs) using BiG-SCAPE^21^. Even though these GCFs originate from only seven metagenomes, they are at similar scale to the diversity of GCFs recorded from 1,038 ocean metagenomes (6,907 GCFs)^19^, 12 antarctica soil samples (2,739 GCFs)^36^, 412 rumen metagenomes (4,565 GCFs)^37^ and 70 metagenomes (2,938 GCFs) from agricultural soils in China ^38^.

BGCs in the peatland microbiome encompassed a wide range of product classes, dominated by ribosomally synthesized and post-translationally modified peptides (RiPPs; n = 1,361), nonribosomal peptide synthetases (NRPS; n = 1,160), terpenes (n = 620), and polyketide synthases (PKS; n = 731), including 149 NRPS-PKS hybrids (**Fig 2; Supplementary table 5**). Within the RiPP category, the most abundant subtypes were RiPP-like clusters (531), RRE-containing clusters (333), ranthipeptides (194), lassopeptides (87), redox cofactors (52), linear azol(in)e-containing peptides (LAP; 39), thioamitides (30), and class IV lanthipeptides (73). Most RiPP subclasses are typically associated with terrestrial bacteria, although LAPs, lassopeptides, and lanthipeptides also occur in marine and freshwater taxa^39^.

**Fig. 2.**
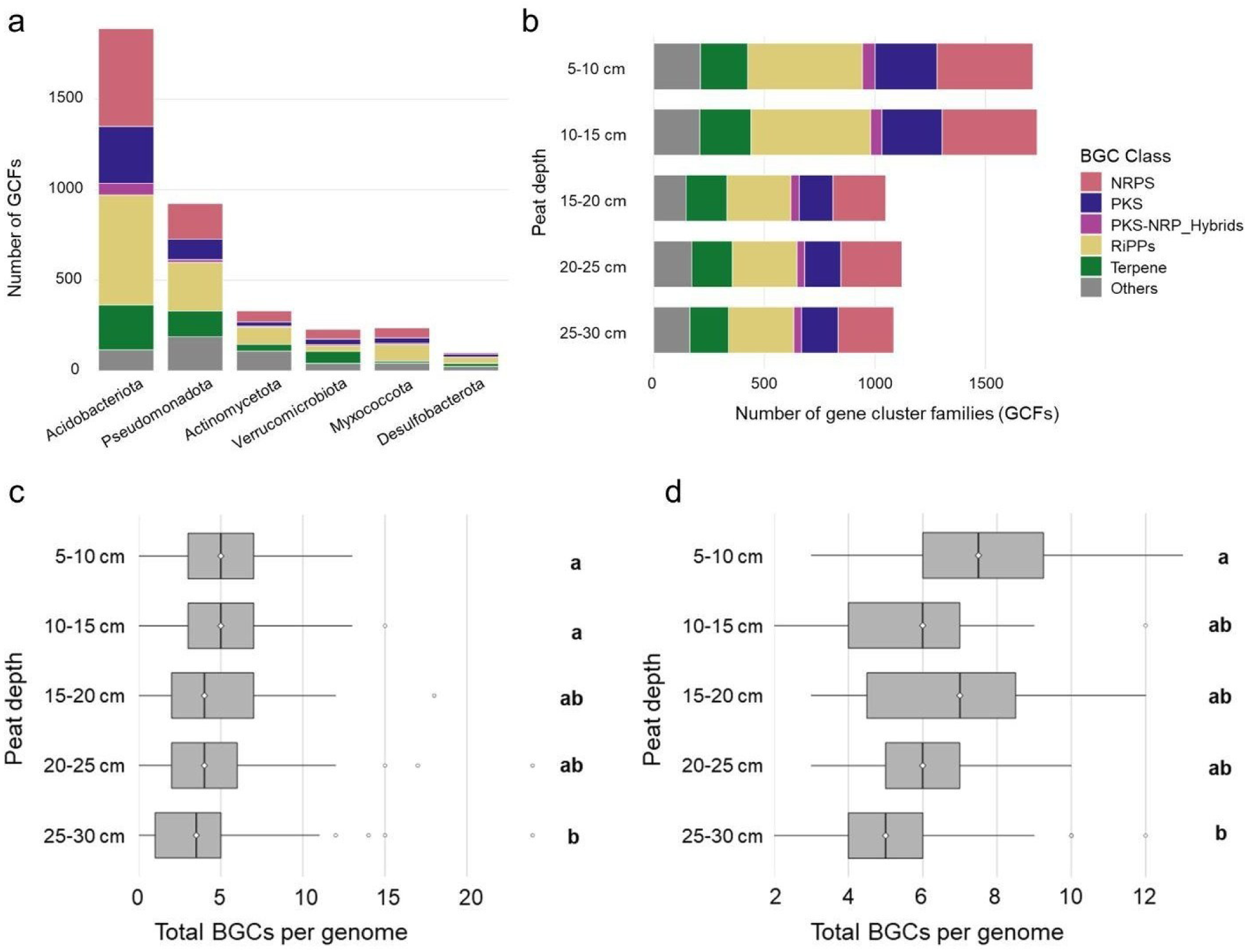
Diversity of Biosynthetic Gene Clusters. **A.** Distribution of BGC classes and their grouping into gene cluster families (GCFs). GCF groupings have been obtained through clustering 9,694 BGCs into families using BiG-SCAPE. GCF abundances are shown per taxa and peatland depth. **B** The mean number of BGCs per genome across all sampled depths are also shown. **C** The number of BGCs per MAG across all depths and **(D)** the mean number of BGCs per MAG belonging to the Acidobacteriota

Gene-cluster families (GCFs) were taxonomically widespread, dominated by Acidobacteriota (1,740) and Pseudomonadota (1,138), followed by Actinomycetota, Verrucomicrobiota (258), Myxococcota (212), and Bacteroidota (122). Additional GCFs were contributed by Desulfobacterota (172), Planctomycetota (77), Nitrospirota (66), and Chloroflexota (38) **(Fig 2)**. Archaeal GCFs were also present (n = 40), with roughly half (21) being of Thermoproteota origin.

MAGs reconstructed from deeper peat layers typically had lower numbers of BGCs than those from upper layers **(Fig. 2C)**. This effect likely reflects the lower biosynthetic capacity of anaerobic taxa, since genomes predicted as anaerobes harbored significantly fewer BGCs per genome than predicted aerobes (Welch t-test, p = 6.56e-08, **Supplementary Fig. 3**). Accordingly, MAGs obtained from the deepest peat layer we sampled often lacked BGCs entirely, while nearly all MAGs from upper layers encoded at least one (**Supplementary Fig. 4**).

BGC-associated regulatory genes differed markedly between the two dominant BGC-encoding phyla. Using Fisher’s exact tests on regulator-type counts, several transcription factor families were enriched in Pseudomonadota BGCs (including LysR, LuxR, GntR, AraC and TetR), whereas Acidobacteriota BGCs showed enrichment of ECF sigma factors (σ24), σ54-associated regulators, and ArsR-family regulators (BH-FDR < 0.05; **Supplementary table 6; Supplementary Fig. 5**)

### Diversity of peatland BGCs

To assess biosynthetic novelty, we compared the 9,694 peatland BGCs to the BGC Atlas^40^ using BiG-SLiCE v2^41^. This approach supersedes earlier comparisons against the BiG-FAM database^42^, which relied on a BIRCH-based clustering scheme in BiG-SLiCE v1 that is less accurate for shorter BGCs^19,41,43^. Only 238 BGCs (2.5%) clustered into existing gene cluster families in BGC-Atlas^40^, which currently comprises nearly two million BGCs from more than 35,000 metagenomic datasets^40^, whereas the remaining 9,456 (97.5%) did not group with any BGC-Atlas^40^ family and thus lacked close relatives in this global reference **(Supplementary Fig. 6**). We also compared our BGCs to experimentally validated clusters in the MiBIG v4.0^44^ database, which further underscored this novelty, with only 10 BGCs (0.1%) showing close similarity (distance ≤ 0.4), including matches to Michiganin A, LanII-2D and SRO15-3108 (class II lanthipeptides), microcin B17, phazolicin and kawaguchipeptin A. The remaining 9,684 had distances > 0.4 to BGCs in the MiBIG database. Thus, virtually the entire peatland BGC repertoire lies outside both genome-derived and experimentally characterized BGC families, highlighting peatlands as a major reservoir of unexplored biosynthetic diversity.

### Peatland MAGs enriched in BGCs

A subset of peatland MAGs stood out as particularly rich in BGCs, with 52 MAGs encoding five or more BGCs (**Fig. 3**, **Supplementary data 3**). The highest counts were from MAGs assigned to Pseudomonadota (formerly Proteobacteria) and Bdellovibrionota. Within Pseudomonadota, two Alphaproteobacteria MAGs each encoded 24 BGCs, including five PKS, five NRPS, and five RiPP clusters. Additionally, 13 other Pseudomonadota (from orders Acetobacterales, Steroidobacterales, Rhizobiales, and Dongiales) carried between 10 and 15 BGCs. We define these MAGs encoding more than 10 BGCs as “talented producers”.

**Fig. 3.**
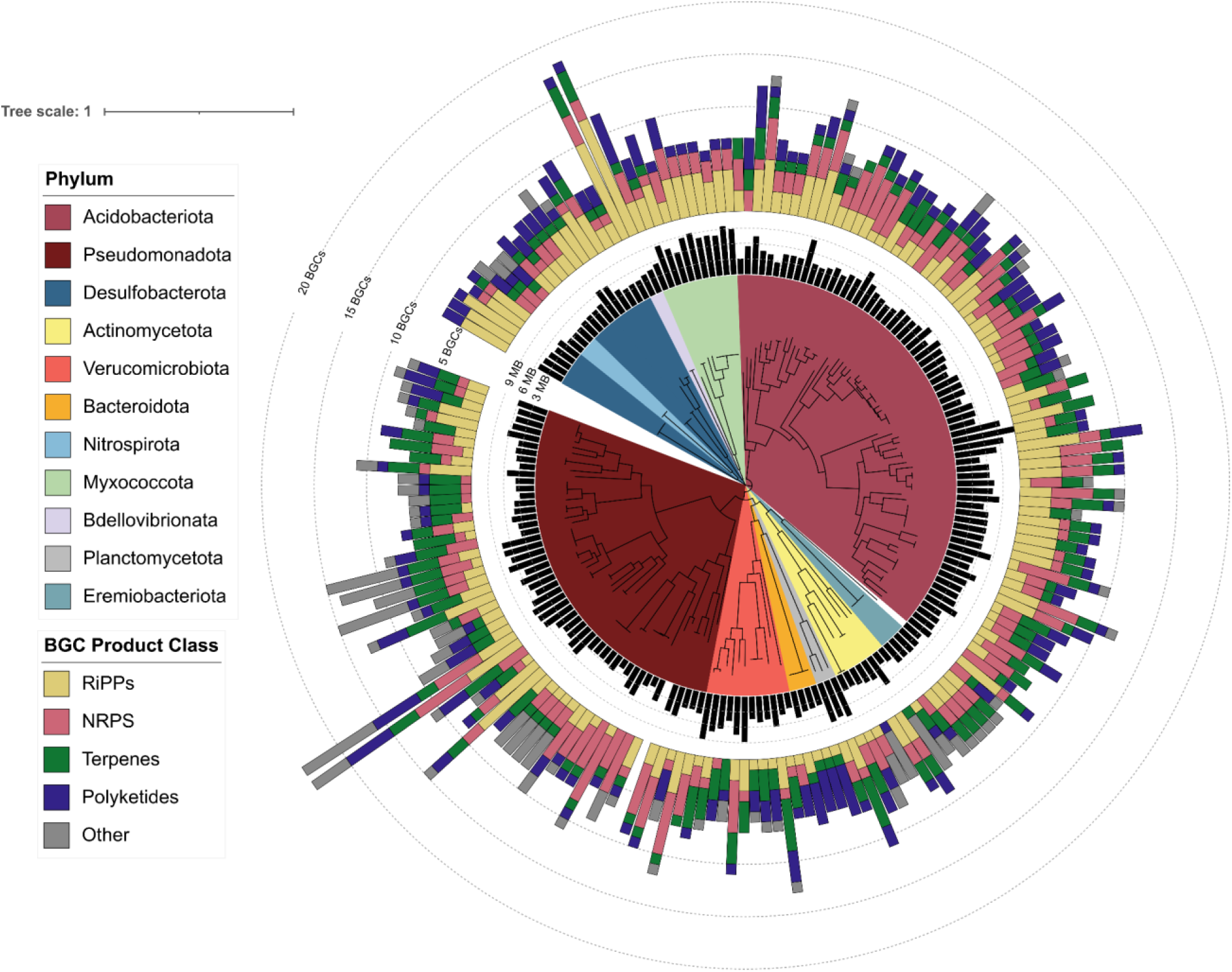
Metagenome-assembled genomes enriched in Biosynthetic Gene Clusters. BGC-rich lineages were identified based on antiSMASH predicted BGCs using recovered MAGs as input. Genomes with at least 5 BGCs are shown in the phylogenetic tree. Phyla are indicated as shaded clusters, genome sizes (Mbp) are shown in the inner barplot, outer barplots show the number of respective BiG-SCAPE categorized classes of BGCs.

Two Bdellovibrionota MAGs (family UBA1018) had 17 and 18 BGCs, of which 10 and 12 were RiPPs, respectively. Fourteen Acidobacteriota MAGs had between 10 and 15 BGCs per genome, which were predominantly affiliated with the Terriglobia and Bryobacteriaceae. Additional BGC-rich genomes included three Verrucomicrobiota, two Myxococcota, and one Desulfobacterota_B (all > 10 BGCs). Among MAGs with fewer than 10 BGCs per genome, notable cases were a Desulfobacterota MAG from the family Syntrophobacteraceae with five RiPP clusters, five different Acidobacteriota MAGs with five RiPPs each, and nine Pseudomonadota and three Acidobacteriota with more than 5 NRPS clusters.

BGCs were also identified in 32 archaeal MAGs, which included a Halobacteriota MAG with four BGCs, including three NRPS clusters, a Thermoproteota and Thermoplasmatota with three BGCs each, and three Halobacteriota had at least two RiPPs. Among these Archaeal MAGs, Thermoproteota also encoded Darobactin, several hydrogen cyanide clusters and RRE-containing clusters, while the Thermoplasmatota also encoded RRE-containing and hydrogen-cyanide clusters.

### *In situ* BGC expression

We quantified BGC expression at contig and genome levels by mapping metatranscriptomic reads to contigs and MAGs and calculating transcripts per million (TPM) for biosynthetic core genes. BGCs were considered expressed when at least one core gene had TPM > 0.25, corresponding to ∼20-40 reads per kilobase. In total, 1,203 BGCs were expressed across the dataset. Per sample, terpenes contributed the largest number of expressed BGCs (mean 129, range 113-167), followed by RiPPs (103, 77-137), with smaller contributions from PKS (36, 26-53) and NRPS (35, 21-61). Acidobacteriota expressed the majority of BGCs (mean 222, 171-287 per sample), followed by Pseudomonadota (40, 14-61). We also summarized the overall investment in BGC expression as the total TPM of core genes. Across depths, Acidobacteriota dominated BGC transcript abundance (mean 205 TPM, 151-346), with smaller but consistent contributions from Pseudomonadota (33.7 TPM, 12.9-66.4), Desulfobacterota (14.9 TPM, 4.4-47) and Verrucomicrobiota (9.9 TPM, 6.44-12.6) (**Fig. 4a,b**).

**Fig 4.**
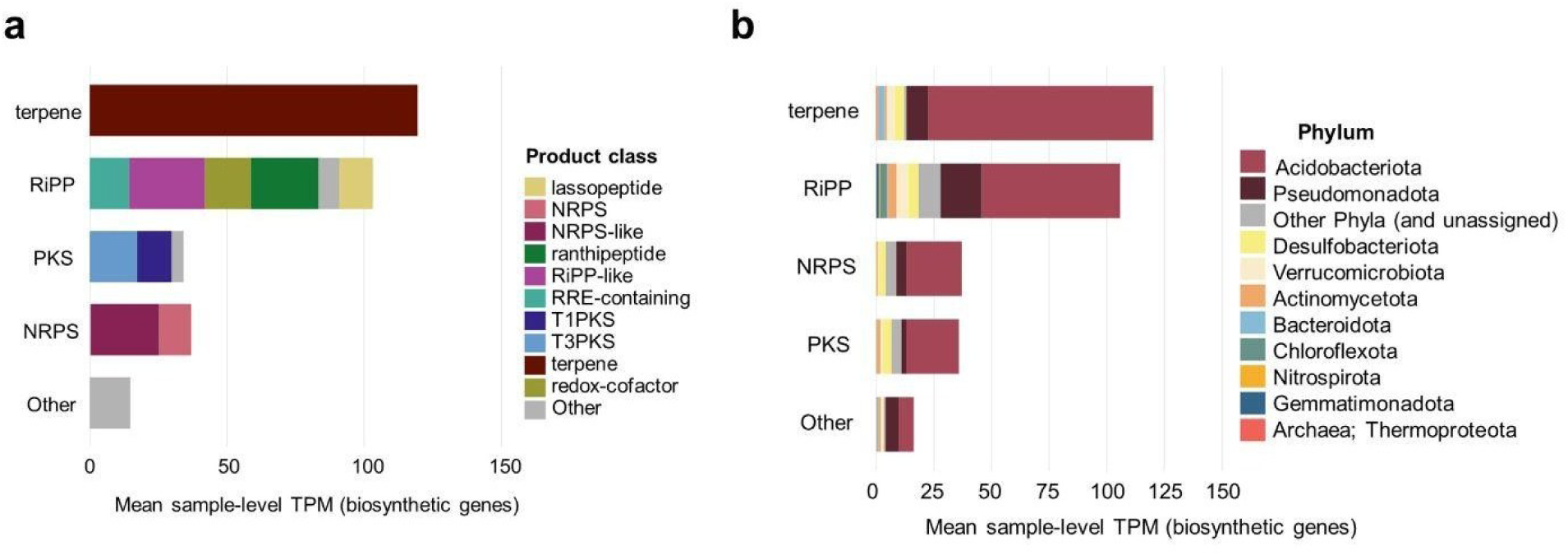
Total expression of core biosynthetic genes from Biosynthetic Gene Clusters. **A.** Expression of BGCs shown as the mean transcripts per million (TPM) per product class per sample and **(B)** the BGC expression by phylum of the producer also shown as TPM values.

### *In situ* BGC expression on genome level

We quantified BGC expression at the genome-level by mapping reads to BGC core genes from active MAGs. MAGs were considered active when the mean TPM values of seven housekeeping genes were higher than 0.5. We defined genome-level biosynthetic potential as the number of BGCs per MAG, from which the proportion of actively expressed BGCs was calculated. Since biosynthetic core genes are essential for metabolite biosynthesis, only those were considered for BGC expression calculations. We observed the expression of accessory BGC genes (linked e.g., to transport, regulation, or unannotated genes) in 801 BGCs in the absence of any reads aligning to biosynthetic core genes.

In our MAG dataset, a total of 281 different BGCs were expressed overall, ranging from 41 - 92 BGCs per sample, or 8.8 - 27% of the biosynthetic potential of the active community **(Supplementary Fig. 6)**. Overall, terpenes accounted for the largest number (67), followed by RiPP-like (43), NRPS-like (29), type III PKS (28), ranthipeptides (28), NRPS (12), RRE-containing clusters (12) and type I PKS (10) **(Fig. 5**). By proportion, ranthipeptides showed the highest expression across samples (33% on average; 18-59% per sample), followed by RiPP-like (24.4%; 13-37%), type I PKS (20.1%; 9-36%), lassopeptides (18.9%; 0-75%), type III PKS (17.3%; 6.9-36%) and NRPS (11.9%; 3.5-27%) **(Fig. 5)**.

**Fig 5.**
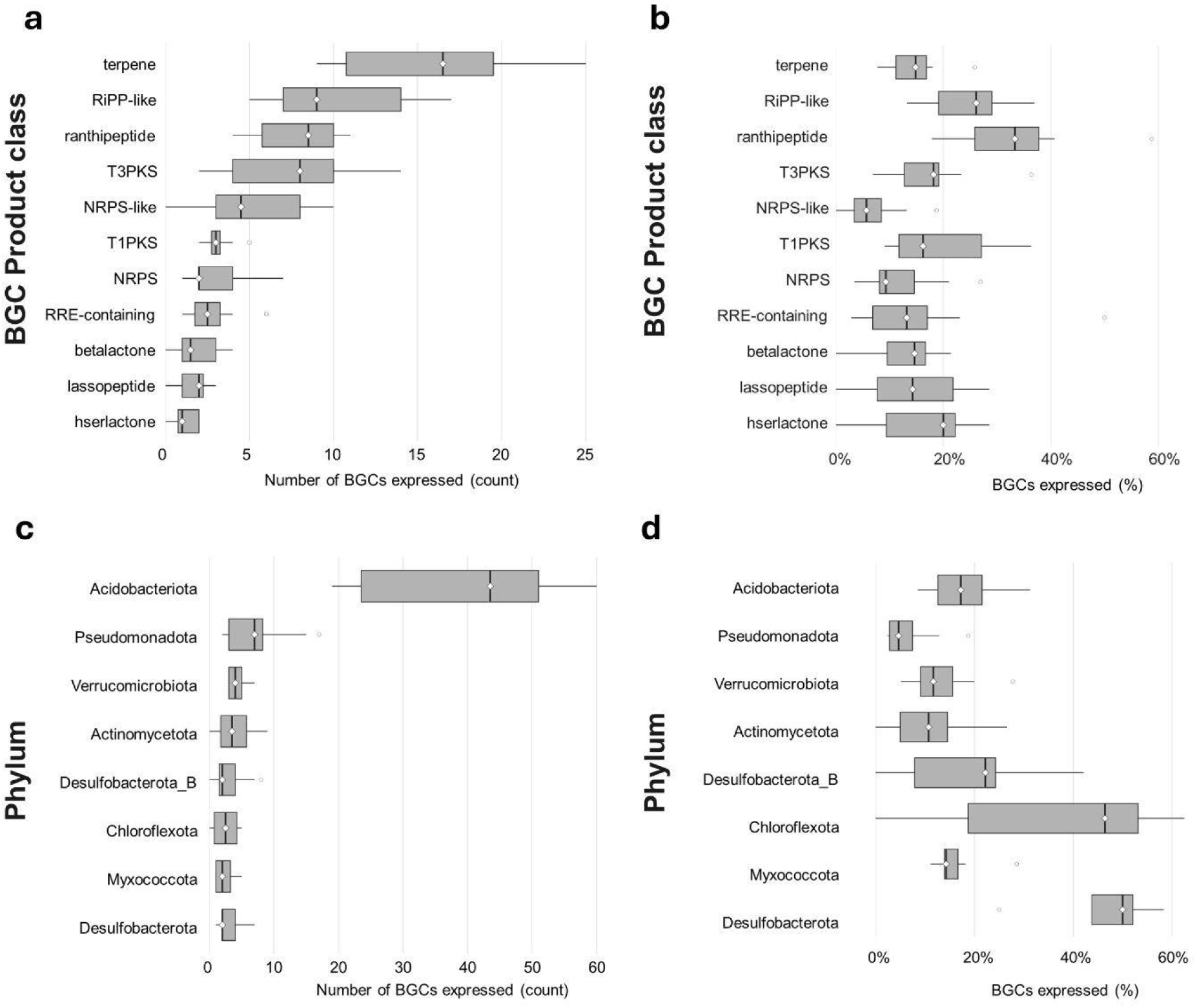
Sample-level expression of biosynthetic gene clusters by product class and phylum. a) Total number of expressed BGCs per sample, grouped by major BGC product classes as assigned by antiSMASH, and (b) proportion of BGCs expressed grouped by predicted product type. c) The number of BGCs expressed per sample assigned to phylum and (d) the proportion of BGCs present in active MAGs that were considered to be expressed per sample.

Acidobacteriota MAGs accounted for most expressed BGCs (n = 163), followed by Pseudomonadota (40), Desulfobacterota (17), Verrucomicrobiota (15) and Myxococcota (8). Acidobacteriota expressed 19-60 BGCs per sample, representing 8.5-31% (mean 21%) of their encoded clusters, whereas Pseudomonadota expressed 2-17 BGCs per sample, corresponding to 2.4-19% (mean 6.4%). Notably, no transcripts were detected from biosynthetic genes in the two most BGC-rich Pseudomonadota MAGs (24 BGCs each; no006_C20-25_concoct.058 and no007_C25-30_metabat2.335) or the two Bdellovibrionota MAGs with 18 BGCs (no003_B15-20_concoct.178 and no006_C20-25_concoct.049). Verrucomicrobiota MAGs expressed 2-7 BGCs per sample (4-8%).

Across all genomes, we observed a clear inverse relationship between BGC repertoire size and the fraction of BGCs that were active (ρ = −0.5, *p* < 0.001, **Supplementary Fig. 9**), indicating that genomes with smaller repertoires tended to express a larger share of their clusters. For Acidobacteriota, this effect was most pronounced below an inflection point of seven BGCs per genome, below which genomes consistently expressed a higher proportion of their biosynthetic capacity than those with larger repertoires. Active Pseudomonadota MAGs, which encoded 3-15 BGCs, expressed no more than five of them in any sample. In contrast, Acidobacteriota encoding 2-13 BGCs routinely expressed a much larger share of their genomic repertoire, including cases of full and near-complete expression. These high expression fractions stand in contrast to the generally low activation rates reported for secondary metabolism in anoxic or energy-limited environments.

### Matching BGC expression to physiological states

We grouped active MAGs based on the expression of genes that are characteristic of different physiological traits, including growth phases, aerobic respiration, stress responses and antimicrobial resistance (AMR) **(Supplementary data 4)**. We examined whether BGC expression is elevated in these MAGs in comparison to BGC expression in all active MAGs combined. Analyses were restricted to MAGs assigned to traits with ≥10 expressed BGCs, with percentages calculated for the fraction of BGCs expressed within a BGC product class for the trait-associated MAG subset versus the overall mean. BGC expression was consistently elevated in genomes expressing markers of aerobic metabolism and stress responses **(Fig. 6; Supplementary data 5)**. For NRPS (overall 0.12), expression was higher with stationary-phase markers (20%, vs 12% overall) and nutrient-stress markers (24% vs 12%), and strongest with AMR genes (50% vs 12% overall). NRPS-like clusters increased with stationary-phase markers (11% vs 6%) and nutrient-stress (17% vs 6%) and were markedly elevated with AMR (23% vs 6%), cytochrome bd quinol oxidase (31% vs 6%), and motility markers (28% vs 6%). Among RiPPs, ranthipeptides increased with stationary-phase (53% vs 31%), nutrient-stress (62% vs 31%), AMR (75% vs 31%), and both bd-type (59% vs. 31%) and aa_₃_-type (49% vs. 31%) terminal oxidases. RiPP-like clusters (overall 22% expressed) were enriched when motility (57%), oxidative-stress (29%), stationary-phase (32%), nutrient-stress (33%), AMR (63%), and aa_₃_-type oxidase (29%) markers were also expressed. For polyketides, T1PKS (overall 17%) was higher with nutrient-stress (38%) and bd-type oxidase (45%). T3PKS (overall 16%) increased with motility (67%), oxidative-stress (24%), stationary-phase (33%), nutrient-stress (43%), AMR (65%), bd-type (66%), and aa_₃_-type oxidase (31%). Terpenes (overall 14%) were elevated with motility (46%), stationary-phase (34%), nutrient-stress (36%), AMR (59%), and with both bd-type (40%) and aa_₃_-type oxidases (25%).

**Fig. 6.**
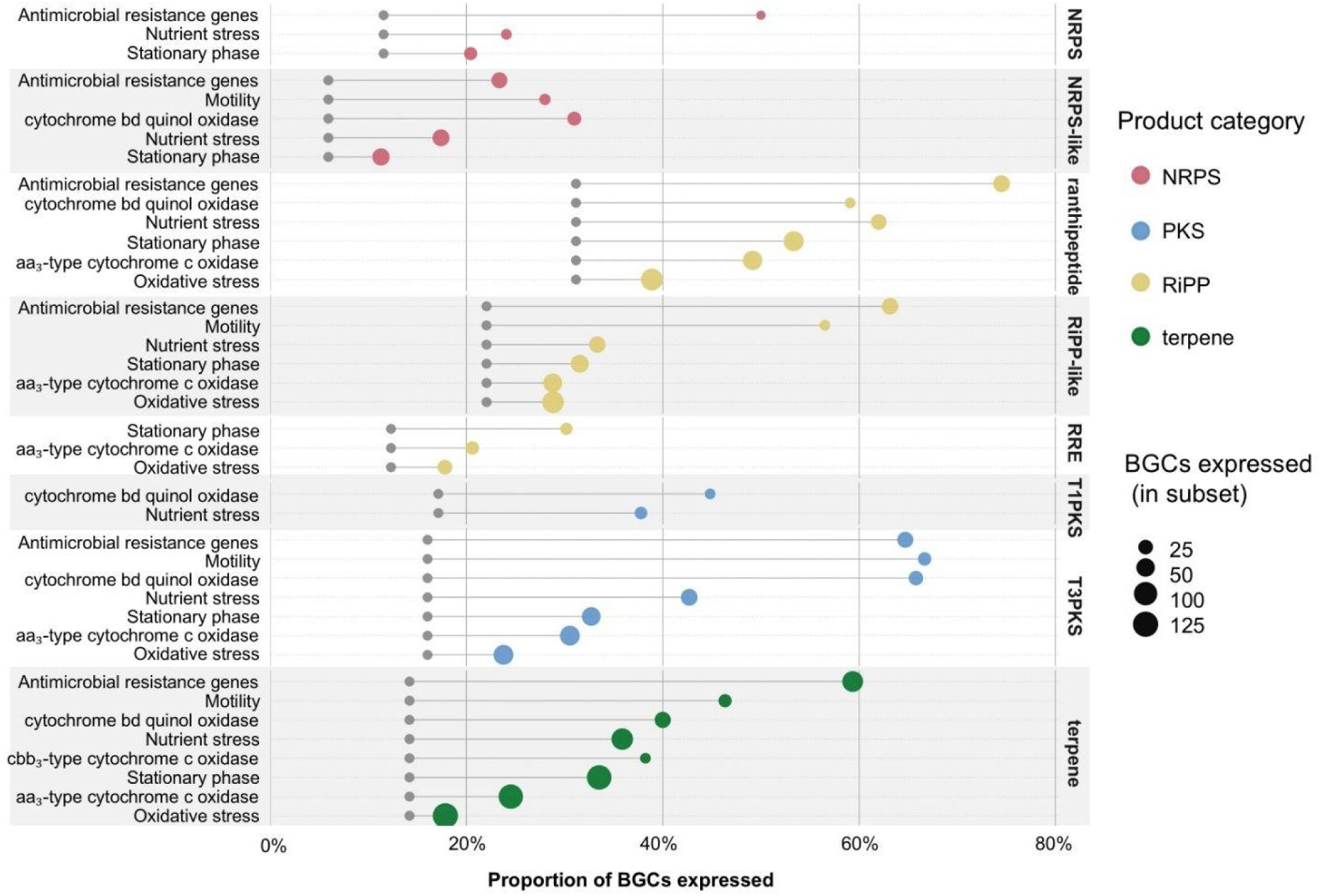
Proportion of Biosynthetic Gene Clusters expressed in MAGs in different physiological states. Expression of BGCs organized by product class (right labels), showing the proportion expressed in a given trait subset (rows), calculated as the proportion of BGCs expressed within that class among MAG by sample observations in which the trait was present. The colored dots represent the trait subset proportion of BGCs expressed and the grey dot on the same row is the overall class baseline across all MAG by sample observations. Point size scales with the number of BGCs expressed in the trait-subset and color shows broader BGC categories. Only trait and product-type combinations with ≥10 expressed BGCs in the subset and a ≥10 percentage-point increase over the class baseline are shown to emphasize substantive effects.

Together, aerobic and high-affinity respiratory systems (aa_₃_ and bd), stationary-phase/nutrient-stress, oxidative stress, motility, and AMR repeatedly coincide with higher BGC expression across NRPS/PKS, terpenes, and multiple RiPP classes, consistent with environmental and physiological control of secondary metabolism *in situ*.

## Discussion

The vast biosynthetic potential of uncultivated microbes is now well established^40,45^, yet how this capacity is realized in natural environments remains poorly resolved. Based on only seven metagenomes, we could show that the studied peatland microbiomes harbor nearly 10,000 BGCs, comparable in scale to global surveys, yet only 8 - 27% were transcribed *in situ*, indicating that environmental and physiological constraints strongly govern secondary metabolism. This selective activation shows that genomic capacity alone poorly predicts realized chemistry and highlights redox state, stress, and metabolic strategy as key determinants of BGC expression. Peatlands thus offer a powerful system for examining how ecological context shapes biosynthetic functions.

Peatland BGCs exhibited an exceptionally high degree of novelty relative to existing genomic and natural product reference databases, reflecting contributions from both classically recognized producers and phylogenetically understudied lineages. These taxa, abundant but historically understudied due to cultivation challenges, harbor some of the richest repertoires of BGCs identified to date, including NRPS, PKS, and RiPP clusters typically associated with well-known producers such as Actinobacteria or Pseudomonadota. Common terrestrial taxa such as Acidobacteriota and Verrucomicrobiota, only recently explored for secondary metabolism^18,20,24^, were major contributors in our dataset, with Acidobacteriota spanning multiple families and metabolic strategies^46,47^ emerging as especially prolific and transcriptionally active producers. Their prominence suggests that broad surveys of Acidobacterial genomes and expression dynamics may reveal substantial, currently overlooked chemical diversity promising for bioprospecting.

We also detected numerous BGCs in anaerobic lineages, such as Chloroflexota, Desulfobacterota, and Planctomycetota, groups recognized as sources of novel biosynthetic capacity in mangrove sediments and anoxic marine systems^48–50^. Their presence underscores that novelty arises not only from classical soil phyla but also from anaerobes adapted to the groundwater-fed, redox-fluctuating conditions of the slightly acidic Schlöppnerbrunnen fen, conditions that support methanogenesis as well as Fe- and sulfate reduction^28,51,52^. Notably, Desulfobacterota, the key sulfate reducers in this fen ^53^, both encoded and actively expressed BGCs, demonstrating that even oxygen-sensitive and energetically constrained guilds contribute to the chemical complexity of peatland microbiomes.

Archaeal lineages, such as Thermoproteota and Halobacteriota, also encoded diverse BGCs, including a darobactin-like RiPP clusters, a group reported to have antibiotic activity against Gram-negative bacteria^54^. The presence of genes associated with hydrogen cyanide biosynthesis in Archaea, known to inhibit cytochrome c oxidases and nitrogenases^55,56^, alongside additional NRPS- and RiPP-like BGCs further indicates that archaeal secondary metabolites may confer competitive or defensive advantages in redox-dynamic habitats.

The coexistence of aerobic decomposers, facultative anaerobes, and strict anaerobes in peatlands creates an unusually heterogeneous biosynthetic landscape. Classical soil taxa and redox specialists jointly expand the known boundaries of secondary metabolism, making peatlands a unique reservoir of chemical novelty shaped by sharp environmental gradients and niche diversification.

In situ BGC expression was dominated by Acidobacteriota, even though the largest biosynthetic repertoires occurred in Alphaproteobacteria and Bdellovibrionota “talented producers”. This decoupling between genomic potential and transcriptional activity reflects fundamental contrasts in life-history strategy. Especially Pseudomonadota are broadly regarded as copiotrophic r-strategists, characterized by genomes enriched in nutrient uptake and transport systems and with high metabolic turnover, motility, and rapid responses to nutrient availability^57–59^. These traits come at substantial energetic cost and are typically favored in resource-rich, oxygenated environments. Thus, their extensive BGC repertoires may therefore represent latent genomic capacity that is deployed only under conditions that meet the regulatory and energetic requirements for activation, conditions that are uncommon or short-lived in peatlands.

In contrast, Acidobacteriota and Verrucomicrobiota are typical oligotrophic, K-selected soil taxa with slower growth, higher carbon-use efficiency, and extensive stress-response machinery, traits that confer stability under chronic resource limitation. Although they encode fewer BGCs than Pseudomonadota, they expressed a much larger fraction of their biosynthetic repertoire, indicating that secondary metabolism is more tightly integrated into their ecological strategy. Many of the expressed Acidobacterial BGC clusters co-occurred with genes for oxidative or nutrient stress responses, and their predicted metabolites, metal chelators, siderophores, and redox-active peptides, are consistent with stress mitigation and resource acquisition.

These ecological differences were evident in regulatory repertoires. Pseudomonadota BGCs were enriched in autoinducer-binding regulators and transcription factors, such as GntR, IclR, AsnC and TetR, which are often associated with quorum sensing, competition and motility^60,61^. Such systems require high cell densities or nutrient cues. In contrast, Acidobacteriota were enriched in ArsR-family regulators and σ^24^ (ECF)- and σ^54^-dependent systems linked to environmental sensing and stress-responsive control^62,63^, which allows coupling between environmental stress and BGC activation. These patterns suggest that copiotrophs primarily activate BGCs in high-resource, competitive settings, whereas oligotrophs deploy smaller but more consistently expressed repertoires as part of a stress-adapted strategy in nutrient-poor settings, such as peatlands.

Despite this selective activation, only a minority of genome-encoded BGCs were expressed across samples. At the genome level, BGC transcription was sparse: the median active MAG expressed only a single BGC, and 52% of active MAGs expressed exactly one. Only a small subset expressed three or more clusters (maximum of seven), and these multi-BGC-active genomes were strongly enriched in Acidobacteriota. This within-genome sparsity mirrors observations from cultured model organisms such as Streptomyces, where up to 90% of BGCs remain silent under standard laboratory conditions (Rutledge & Challis, 2015). Such low activation rates reflect the high metabolic cost of secondary metabolite biosynthesis, which diverts precursors from primary metabolism and relies on multilayered regulatory systems integrating nutrient status, stress responses, and community-level cues^64–66^. In peatlands, these constraints appear to selectively favor Acidobacterial activation of BGCs under environmental stress, while rendering the larger but more energy-demanding Pseudomonadotal repertoires largely dormant.

Activation of PKS, terpene and some RiPP classes were consistently associated with aerobic respiration and oxidative stress, indicating that microbes invest in these energetically demanding pathways when respiratory energy is available and competitive or stress-related cues intensify. Many of the megasynthases depend on O_₂_-dependent tailoring steps^67^, making their expression most feasible under oxic conditions. Similar patterns are well known from model producers such as *Streptomyces*, where most BGCs remain transcriptionally silent under standard laboratory conditions^68^. In our dataset, RiPP expression correlated with antimicrobial-resistance markers and two-component systems, consistent with contact- or stress-induced defence responses. Together, these observations support the view that BGC activation is governed by energetic constraints and cellular stress signaling, rather than constitutive expression.

Despite the strong association between certain BGC classes and aerobic metabolism, expression was detected across all peat depths, including in deeper, more anoxic layers. This suggests that secondary metabolism persists even where biosynthetic capacity is reduced and energy availability is limited. In deeper peat layers, BGCs are likely repurposed toward stress mitigation, such as redox balancing, metal chelation, or protection against oxidative and envelope stress, rather than classical antibiotic competition^50^. Comparable functional shifts toward redox-active or protective secondary metabolites have been reported in anoxic marine sediments and cold seeps^48,49^. The ability of Acidobacteriota to express BGCs in both oxic surface layers and deeper anoxic zones underscores their ecological breadth, enabling them to compete with fast-growing aerobic decomposers as well as slow-growing anaerobic specialists. Sustained BGC activity across depths thus points to a broader ecological function in stress tolerance and metabolic stability under fluctuating oxygen and nutrient conditions.

## Conclusion

Our study shows that cataloguing BGC diversity alone is insufficient to infer ecological function. Although peatland microbes encode nearly 10,000 BGCs across a broad phylogenetic range, only a small fraction is expressed *in situ*, and expression is shaped more by physiological state and environmental conditions than by genomic capacity. Acidobacteriota were the dominant contributors to BGC transcription, even in deeper, energy-limited layers, indicating a central role for stress-and adaptation-related secondary metabolism. Overall, our findings reveal that heterogeneous, redox-structured environments harbor extensive biosynthetic novelty, but only select conditions trigger its activation. BGCs are everywhere, but the environment selects.

## Methods

### Site description and sampling

Peat soil cores were collected from the Schlöppnerbrunnen fen, located in the northern Fichtelgebirge, Northern Bavaria, Germany (50°07′55″N, 11°52′52″E). The fen covers approximately 0.8 ha and is characterized by slightly acidic soils on granite bedrock. Vegetation is dominated by *Carex canescens*, *Carex rostrata*, *Juncus effusus*, *Molinia caerulea*, and *Eriophorum vaginatum*. Peat accumulation varies between 30 and 70 cm depth^51^. The Schlöppnerbrunnen fen is fed by anoxic, Fe(II)-rich groundwater leading to an enrichment of total iron in the upper peat layers. Groundwater levels fluctuate between 10 and 30 cm below the surface, influencing mean oxygen penetration depths, which was previously reported to be up around 16 cm deep^69^. Across depths of 0 - 30 cm, the fen has a pH of ∼ 4.7, total iron content of 105 - 170 µmol.g^-1^ ^28,51^ and sulfate content of 20 to 240 µM^70^. The Schlöppnerbrunnen fen has been the focus of extensive biogeochemical and microbial research^27,28,52,70,71^.

For this study, we analyzed data from two parallel peat cores (B, and C) collected in July 2020 using a soil corer. Cores were sectioned in the field at 5 cm intervals down to 30 cm depth, and each segment was homogenized. Samples were flash frozen, transported on dry ice, and stored at −20 °C until DNA and RNA extraction. Due to the very dry conditions in the field, peat from the upper 0-5 cm depth was not included in the extractions. Despite multiple attempts, RNA could not be recovered from the deeper 20-30 cm layers, likely due to inhibitory compounds and high iron content.

### Nucleic acid extraction and sequencing

Total RNA was extracted using the RNeasy Powersoil Total RNA Kit (Qiagen, Hilden, Germany) following the manufacturer’s instructions. Potentially present genomic DNA was digested with TURBO^Tm^ DNase (ThermoFisher Scientific, Darmstadt, Germany) before being purified using the RNA Clean and Concentrator-5 kit (ZymoResearch, Freiburg, Germany). Purified RNA was quantified through fluorometry, using a Qubit® 3 fluorometer, and RNA HS assay reagents (ThermoFisher Scientific). Purity was assessed using spectrophotometry with a NanoDrop® microvolume spectrophotometer (ThermoFisher). RNA integrity was checked using agarose gel electrophoresis. RNA samples were subjected to Illumina sequencing library preparation using the NEBNext Ultra II Directional RNA Library Prep kit (New England Biolabs, Frankfurt am Main, Germany), aiming for an insert size of ∼200 bp. Finalized libraries were quantified through fluorometry, and their size distribution was checked using a Bioanalyzer® instrument (Agilent Technologies, Waldbronn, Germany) and the RNA 6000 pico kit. Sequencing was carried out at the sequencing core facility of the Leibniz Institute on Aging - Fritz Lipmann Institute on an Illumina NovaSeq device using an S2 flowcell in paired-end mode (2 x 150 bp).

### Sequence data processing

Metagenome reads were trimmed using *TrimGalore* (v0.6.6) ^72^ with a specified -- length of 75 and --quality of 28 to remove low quality end bases and filter out sequences of <75 bp. Forward and reverse reads were merged using *Flash* (v1.2.11) ^73^ with a -- max-overlap of 151 bp. Metagenome sequences were assembled using *SPAdes*^74^ (v3.15.3) in metagenome mode with kmer sizes 21, 33, 55, 77, 99, 111, and 127. Metagenome coverage / redundancy was estimated using Nonpareil v3.5.5^75^. Assemblies >2000 bp were binned using *MetaBat2* (v2.15)^76,77^, *Concoct* (v1.1.0)^78^, and *Maxbin* (v2.2.5)^79^. Bins were then evaluated with *DAS Tool* (v1.1.2) ^80^ to select the best non-overlapping bins. A selection of 573 bins from seven peat samples, had completeness > 70% and contamination < 10% which were used for further analyses (**Supplementary table 3**). Bin quality, percent of binned population, and estimated percent of community were assessed using *CheckM* (v1.2.2) ^81^.

### Annotation

Gene calling was done using Prodigal v2.6.3, as implemented in Anvi’o v8^82,83^, and then annotated using kofams^84^, COG20 and CAZy (dbCAN2) ^85^ databases using the commands *anvi-run-kegg-kofams*, *anvi-run-ncbi-cogs* and *anvi-run-cazymes* respectively. Additional annotations were done for MAGs using METABOLIC v4.0, using the <u>METABOLIC-G.pl</u> script^86^. Gene calls were exported from Anvi’o contig databases using the anvi-get-sequences-for-gene-calls with the –export-gff3 setting. Contig-level taxonomy was determined using MMseqs2 ^87^, using GTDB Release 220^88^ as reference. MAG taxonomic annotation was performed using GTDB-tk^89^, using GTDB Release 220.

### BGC identification

Biosynthetic gene clusters (BGCs) were identified from metagenome contigs longer than 5 kb using *AntiSMASH*^35^ (v7.0.1) with the settings –fullhmmer –pfam2go – cc-mibig –cb-subclusters –rre –smcog-trees –asb –cb-knownclusters. The BGCs identified within each sample were further clustered into biosynthetic gene cluster families (GCFs) as a means to dereplicate them using *BiG-SCAPE* (v1.1.9) ^21^ with settings --include_singletons --mibig --mode auto and 0.3 similarity threshold. Clustering in *BiG-SCAPE* is done through the calculation of pairwise distances between BGCs using a weighted combination of the Jaccard, Adjacency Index, and Domain Sequence Similarity indices. Furthermore, BiG-SCAPE classifies BGCs into terpenes, NRPS, RiPPs, PKSother, PKS type I, NRP-PKS hybrids, and others, and these groups are hereafter referred to as *BiG-SCAPE* classes. BGC abundance in metagenomes were calculated by alignment of previously quality-controlled metagenome reads against the seven metagenome assemblies using *Bowtie2*^90^ (v2.4.5). BGC novelty was assessed by comparing GCFs against the BGC-Atlas^40^ and MiBIG 4.0^44^ databases using *BiG-SLiCE* 2.0^41^. BGC-Atlas comprises more than two million BGCs from various metagenomic sources, while the MiBIG database contains all experimentally validated BGCs.

### Metatranscriptomics

Raw metatranscriptome reads were quality filtered and sequencing adapters removed using *fastp*^91^ (v0.23.4) with default settings, followed by rRNA filtering using Ribodetector^92^ (v0.3.1), using default settings. For calculation of total BGC transcription, ribosomal RNA-depleted QC-reads were aligned to whole assemblies using bowtie2 (v2.4.5).

For MAG-level analyses, ribosomal RNA-depleted, quality-filtered reads were aligned to a dereplicated set of MAGs (n = 369; 98% average nucleotide identity, dRep^93^ v2.6.2, --S_ani 0.98). Read mapping was performed with Bowtie2 (v2.4.5) using default settings, and SAM files were converted to BAM with samtools^94^ (v1.7). Aligned BAM files were then filtered with reformat.sh from BBTools^95^ (v39.01) using pairedonly=t, primaryonly=t and idfilter=0.95, and converted to gene counts with featureCounts^96^ using -s 0 -t CDS -p --countReadPairs. In total, 131 million read pairs mapped to MAGs (4.5-20.8 million per sample). Counts < 5 per sample and < 20 across all samples were removed, and remaining counts were normalized to transcripts per million (TPM) in R.

BGC expression was assessed using different criteria in the contig-level and MAG-level analyses. In the contig-level dataset, a BGC was classified as expressed if at least one biosynthetic core gene had TPM > 0.25 (approximately 20–40 reads per kilobase). For MAG-level analyses, expression was evaluated only in “active” MAGs, defined as genomes with a mean TPM > 0.5 across seven single-copy housekeeping genes: rpoB (K03043), rpoC (K03046), gyrA (K02470), tuf (K02358), infB (K02519), atpD (K02112), and sigA (K03086). BGCs were considered expressed when core biosynthetic genes, as identified by antiSMASH v7.0^35^, met at least one of the following criteria: (i) mean biosynthetic core-gene TPM ≥ 75% of the mean TPM of the housekeeping genes, (ii) any core gene with TPM greater than the genome-wide housekeeping-gene mean, or (iii) biosynthetic core gene TPM > 0.8, corresponding approximately to 20–30 reads per kilobase. Multi-mapping of metatranscriptomic reads was minimized by dereplicating MAGs at 98% ANI prior to mapping.

Across samples, the mean number of active MAGs was 82 (range 45-119 per sample) from the 349 MAGs used for BGC expression mapping. These active MAGs contained 282-680 BGCs per sample, corresponding to 969 distinct BGCs across all samples, which we refer to as the total biosynthetic potential. Of these, 803 BGCs were detected as expressed at least once, representing 281 unique BGCs.

### Data analyses

All statistical and associated data analyses were performed in R version 4.3.0 ^97^, unless stated otherwise. R packages *dplyr* (v1.1.2)^98^, *forcats* (v1.0.0)^99^ and *stringr* (v1.5.0)^100^ were used for processing large data tables, and *ggplot2* (v3.5.2)^101^ was used to generate all figures, except for Fig. 3 which was created using iTol^102^.

Differences in biosynthetic capacity between groups of MAGs were assessed using two-sided Welch’s t-tests using the *t.test* function in base R^97^ (v4.3.0). Genomes predicted as anaerobes and aerobes were compared for the number of BGCs per genome. The relationship between BGC repertoire size and the fraction of BGCs expressed per MAG was quantified using Spearman’s rank correlation, using the *cor.test* function in base R^97^ (v4.3.0).

The enrichment in regulatory domains in BGCs from the Acidobacteriota and Pseudomonadota was assessed using Fisher’s exact tests on tables of SMCOG-annotated regulators extracted from antiSMASH BGC annotations, and compared using the *fisher.test* and *p.adjust* commands in base R^97^ (v4.3.0).

Associations between BGC expression and physiological traits were evaluated by grouping MAG x sample observations according to the expression of marker genes for aerobic respiration, growth phase, stress responses and antimicrobial resistance **(Supplementary data 4)**. For each trait and BGC product class, we compared the proportion of expressed BGCs in the trait-associated subset to all remaining observations using two-sided Fisher’s exact tests. Multiple testing was controlled within each BGC class using the Benjamini-Hochberg false discovery rate (FDR). These analyses were performed using the *fisher.test* and *p.adjust* functions in base R (v4.3.0)^97^.

## Data availability

Raw sequencing data generated in this study is available in the NCBI Sequence Read Archive under BioProject PRJNAxxxxxx. Metagenome assembled genomes (MAGs) and Biosynthetic gene clusters generated in this study have been deposited in Zenodo under accession 10.5281/zenodo.xxxxxxx.

## Code availability

Code used for data analysis and to generate figures have been deposited in Zenodo under accession 10.5281/zenodo.xxxxxxxx.

## Acknowledgements

This research was supported through the Collaborative Research Center Chemical Mediators in Complex Biosystems (SFB 1127 ChemBioSys, Project Number 239748522), which is funded by the Deutsche Forschungsgemeinschaft (DFG) and was awarded to KK and CEW. Additional funding was provided by the Geobiology and Low Temperature Geochemistry program at the National Science Foundation (EAR-1833525). The authors thank Pierre Stallforth and Anan Ibrahim for their helpful comments on this research.

## Author contributions

KK, CEW, ZRH, CC and RH designed the study, KK and CEW collected and processed field samples. CC, RH, CEW and ZRH performed initial bioinformatic analyses and ZRH and CEW performed analyses on biosynthetic gene clusters and KK, CEW and ZRH wrote and revised the manuscript.

## Competing interests

The authors declare no competing interests.

## Notes

### Competing Interest Statement

The authors have declared no competing interest.

